# Dietary emulsifiers and host inflammation synergistically drive genomic evolution of Crohn’s disease-associated *E. coli* toward enhanced pathogenicity

**DOI:** 10.64898/2026.04.20.719593

**Authors:** Héloïse Rytter, Caroline Chevarin, Lucas Martin, Emma Bruder, Jérémy Denizot, Olivier Tenaillon, Olivier Espéli, Aurélien Birer, Emilie Viennois, Nicolas Barnich, Benoit Chassaing

## Abstract

**Background and Aims:** The rising incidence of Crohn’s disease (CD) in Westernized countries has been linked to changes in diet and increased consumption of food additives, yet the mechanisms by which these factors fuel intestinal inflammation remain unclear. Adherent-invasive *Escherichia coli* (AIEC), a pathobiont involved in CD pathogenesis, lacks a clear genetic hallmark but exhibits intestinal colonization and virulence traits, raising questions about the evolutionary forces promoting its emergence among select individuals. Here, we investigated how chronic exposure to two common dietary emulsifiers, carboxymethylcellulose (CMC) and polysorbate 80 (P80), along with host inflammation, drives AIEC genomic evolution and pathogenic potential.

**Methods:** Wild-type and Il10-deficient mice were monocolonized with AIEC and chronically exposed to CMC, P80, or water. Bacterial isolates were collected and analyzed for genomic diversification, mutations, and phenotype both *in vitro* and *in vivo*.

**Results:** Emulsifiers accelerated AIEC genomic diversification and selected for mutations linked to increased motility, invasion, and pro-inflammatory activity. Moreover, dietary emulsifier-evolved strains displayed a marked fitness advantage *in vivo*, outcompeting their counterparts in murine hosts, with the greatest advantage observed when evolution occurred under inflammatory conditions. Notably, evolutionary pathways and phenotypic outcomes were shaped by both emulsifier and the host’s inflammatory status, highlighting synergy between diet and host genetics in fostering pro-inflammatory pathobionts.

**Conclusion:** These findings provide an evolutionary framework connecting modern dietary habits to the emergence of pathogenic AIEC strains, and underscore the importance of dietary interventions in individuals at risk for inflammatory bowel disease.

## Introduction

Throughout human history, dietary habits have evolved under the influence of biological, environmental, technological, and sociocultural forces. Industrialization marked a decisive shift, introducing food additives designed to extend shelf life, enhance texture, and support large-scale production, particularly across Western countries^1^. While these innovations have diversified our food supply, the rapid emergence of novel dietary components, especially food additives, raises important questions about human physiological and microbial adaptation. The mismatch between our ancestral diets and modern eating patterns may underlie the growing prevalence of noncommunicable diseases, such as Crohn’s disease (CD), observed in Western societies^2–4^.

Although CD lacks a single microbial signature, it is frequently associated with a dysbiosis characterized by a decline in beneficial bacterial phyla and a proliferation of mucosa-associated *Enterobacteriaceae*, most notably adherent-invasive *Escherichia coli* (AIEC)^5–9^. AIEC strains are characterized by enhanced adherence to and invasion of intestinal epithelial cells, features suspected to promote chronic mucosal inflammation^10–12^. However, unlike classical pathogenic *E. coli*, AIEC strains do not share a unique genetic signature, and are phylogenetically diverse, suggesting their independent emergence in response to diverse selective pressures. Indeed, studies tracking AIEC evolution in animal models have reported that host-associated factors, including chronic inflammation, can drive the selection of more virulent lineages^13,14^.

Among environmental factors, diet is a major determinant of gut microbiota composition and host inflammatory status^15,16^. Epidemiological studies have associated high consumption of ultra-processed foods and their commonly added dietary emulsifier, including carboxymethylcellulose (CMC) and polysorbate 80 (P80), with increased risk of CD^4,17^. These emulsifiers have been shown in mouse models to disrupt the gut microbial ecosystem and potentiate low-grade intestinal inflammation^18,19^. Moreover, both human and animal studies indicate that the detrimental impact of dietary emulsifiers depends on resident microbiota^20,21^.

Our previous work has identified AIEC as a key direct target of emulsifier-induced dysbiosis and inflammation. Indeed, colonization with AIEC was observed to be sufficient to make mice highly susceptible to the harmful effects of CMC and P80, and direct exposure to emulsifier drives the expression of AIEC’s virulence genes^22^. However, the long-term consequences of chronic emulsifier exposure on AIEC evolution remain unknown. We hypothesize here that dietary emulsifier consumption, either alone or in the context of host inflammation, could serve as a selective pressure driving AIEC adaptation to favor its ability to stably colonize the gastrointestinal tract, together with increasing its pathogenic potential. To address this, we tracked the evolution of AIEC in monocolonized wild-type (WT) and *Il10*-deficient mice subjected to prolonged CMC or P80 exposure, and assessed the resulting bacterial genetic and phenotypic changes. Our findings importantly revealed that dietary emulsifiers are sufficient to drive genomic and phenotypic evolution of AIEC bacteria, ultimately leading to their increased pathogenic potential. Altogether, these data provide new insight into how modern dietary practices might have contributed to the emergence of pathobionts in Crohn’s disease.

## Materials and Methods

### Mice monocolonization experiment

As previously reported in Viennois *et al* ^22^, C57BL/6 inbred mice (C57BL/6NTac) and Germ-free C57BL/6 *Il10*^-/-^ inbred male mice (C57BL/6NTac-*Il10^em8Tac^*; Taconic model GF-16006) were maintained in isolated ventilated cages Isocages (Techniplast, West Chester, PA, USA^23^) to prevent exogenous bacterial contamination. All mice were bred and housed at Georgia State University (Atlanta, GA, USA) under institutionally-approved protocols (IACUC # A17047 and A18006). Mice were fed autoclaved LabDiet rodent chow # 5021. Mice used in this study were 4-5 weeks old, at which point AIEC reference strain LF82 was administered in drinking water. AIEC LF82 was grown overnight in 200 mL of LB at 37°C without agitation and bacterial suspensions with a 620nm optical density (OD) of 2.0 were placed in the water bottles of germ-free mice. One week later, water solution was put back and supplemented with CMC or P80 (1.0%). After 12 weeks, mice were euthanized and their feces and ceacum were collected for downstream analysis (**Figure 1A**).

**Figure 1.**
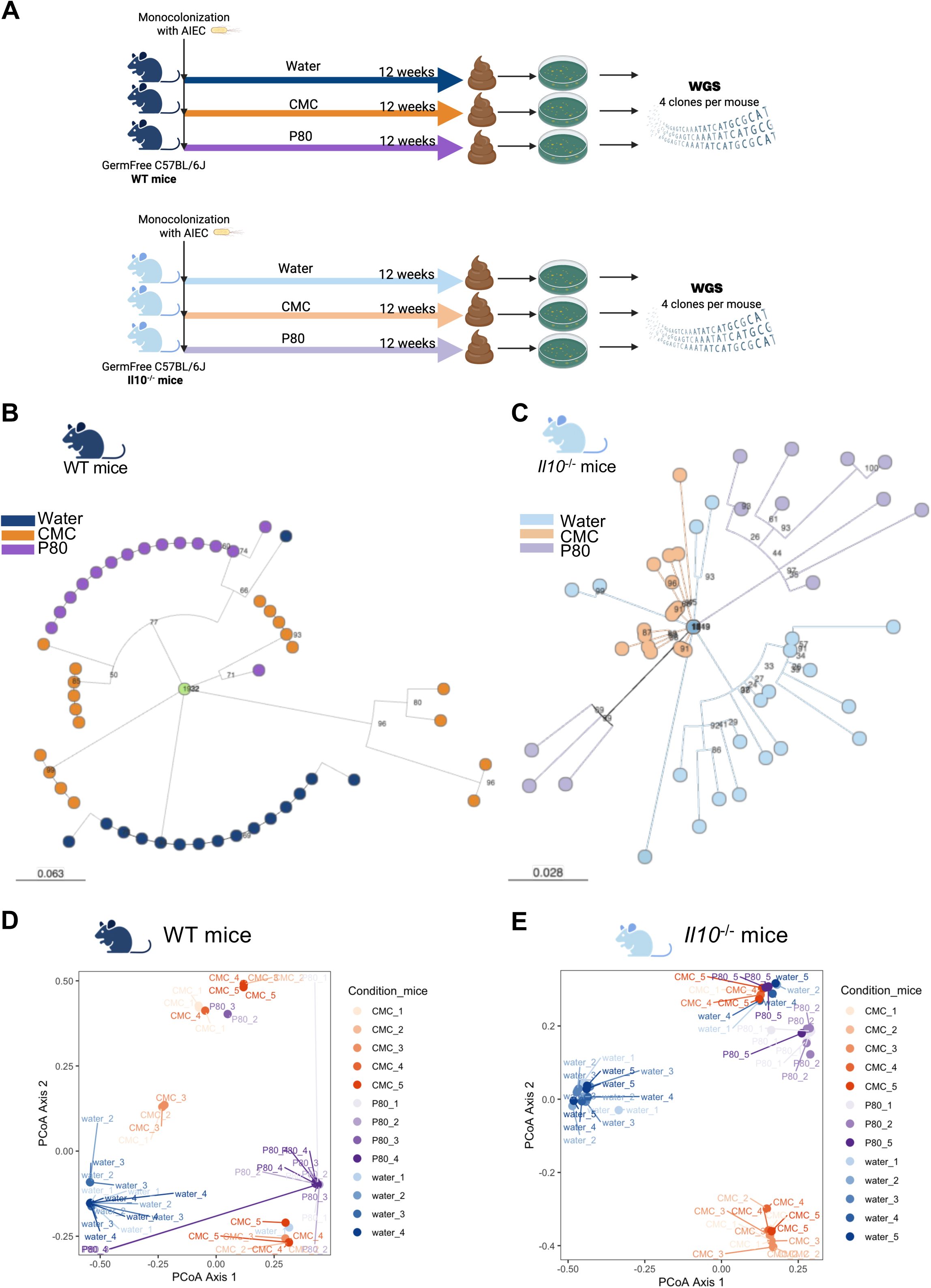
Host genetic background and dietary emulsifiers shape AIEC genomic evolution. (**A**) Overview of the experimental plan used. Four to five WT and *Il10*^-/-^ mice were mono-colonized with AIEC reference strain LF82, and supplied with drinking water containing CMC, P80 or no additive as control for 12 weeks. After euthanasia, AIEC clones were recovered from fecal pellets (WT mice) or cecal contents (*Il10*^-/-^ mice) on Drigalski agar plates. Four colonies per mouse were randomly picked for whole-genome sequencing. (**B-C**) Maximum⍰likelihood phylogenies of the evolved AIEC isolates, constructed from the set of SNPs identified relative to the LF82 reference genome. Such analysis was performed on strains from WT (**B**) and *Il10*^-/-^ (**C**) mice. Branch colors indicate treatment group and bootstrap values are shown on each node. (**D-E**) PCoA of the mutational profiles of each strain isolated from WT (**D**) and *Il10*^-/-^ (**E**) mice. Numbers indicate the mice number

**Figure 2.**
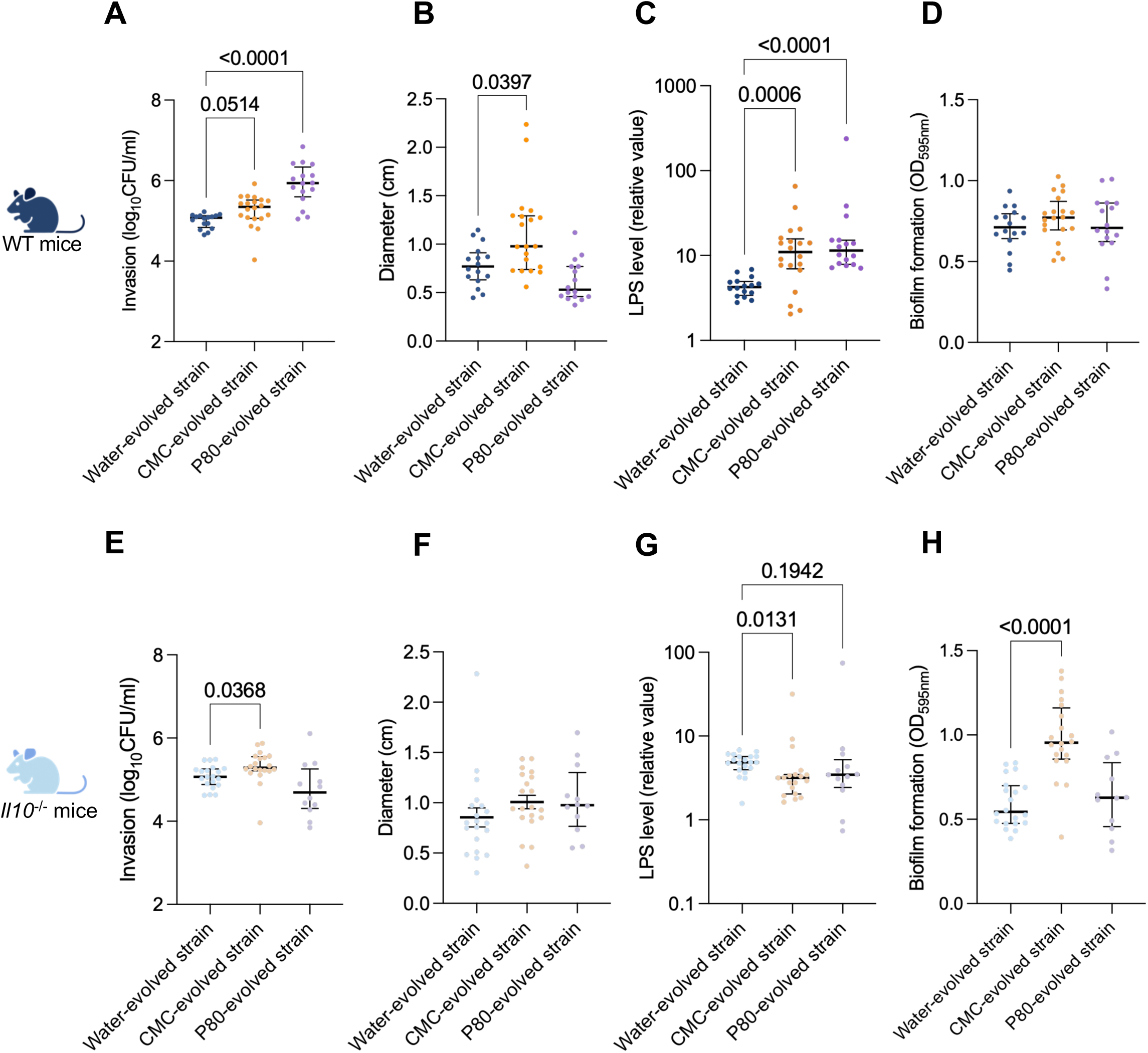
Dietary emulsifiers-driven genomic evolution of AIEC increase their pathogenic potential *in vitro.* *in vitro* pathogenic traits of isolates from WT (**A-D**) and *Il10*^-/-^mice (**E-H**) were assessed by measuring invasiveness of intestinal epithelial cells (**A, E**), swimming motility (**B, F**), bioactive LPS level normalized by OD_600nm_ (**C, G**) and biofilm formation (**D, H**). Data are shown as the median with interquartile range. ∗p < 0.05 compared to water-isolated strains, determined by Kruskall-Wallis test.

### AIEC isolation from mono-colonized mice

To analyze the impact of dietary emulsifier on AIEC evolution, ceca or feces from AIEC LF82-monocolonized C57BL/6 and C57BL/6 *Il10*^-/-^ mice (N= 4-5) were homogenized and plated on selective Drigalski media to recover LF82 clones. For each mouse, four colonies were randomly picked to establish a library of evolved isolates (**Figure 1A**).

### Bacterial Strains and Culture Conditions

For growth analysis, AIEC strains were cultured in LB broth at 37°C without shaking. OD at 600 nm was measured every 30 minutes (min) using a SpectraMax ABS Reader.

#### Construction of kanamycin resistant strains

Ancestral LF82-Km^R^, CMC evolved-strain-Km^R^, P80 evolved strain-Km^R^ both from WT mice and water evolved strain-Km^R^ strain from *Il10^-/-^* mice were constructed employing the mini-Tn*7* system in bacteria with single *att*Tn*7* sites. As previously described^24^, GFP tagged-kanamycin resistance gene was cloned using Gibson method^25^ into an appropriate mini-Tn*7* vector to generate pUC18R6KT-mini-Tn7T-Km-*gfp*. The mobilizable mini-Tn7-based vector was delivered by conjugation into the concerned *E. coli* strains using *E. coli* MFDpir46 and the helper *E. coli* MFDpir/pTNS3 strain encoding the *tnsABCD* genes necessary for the transposition of mini-Tn7 at the *attTn7* insertion site^26^. Selection of insertion-containing strains was followed by PCR verification.

### Whole genome sequencing of evolved AIEC clones

Total DNA were extracted from the isolated AIEC clones using the Wizard® HMW DNA Extraction Kit (Promega), following the manufacturer’s instructions. Genomic sequencing was performed on Illumina Nextseq 2000 (QC ADN Illumina DNA prep 200 cycles) sequencing runs at the Institut Cochin (Paris, France). Bioinformatic analysis were performed as previously described^14^. Briefly, raw reads were filtered and trimmed to ensure good quality using Cutadapt (version 3.2)^27^. Any pair containing Ns, homopolymers (10 nucleotides or more), or those longer than 151 bp were discarded. Sequences were trimmed with a quality cutoff of 25 at both ends for both reads, Illumina adapters were removed, and sequences shorter than 50 bp were discarded. A filter was applied to remove any adapter leftovers using Trimmomatic^28^ and sequences shorter than 105 bp were discarded. Mutations were identified using the Breseq tool ^29^ by comparing the obtained genomes with the reference genome of *E. coli* LF82 (PRJNA33825) using default options (i.e. mutations identified by a threshold frequency□>□0.05) and by subtracting the mutations detected in original glycerol stock of the LF82 clones used to inoculate germfree mice.

### Construction of phylogenetic tree

Single-nucleotide polymorphism (SNP) discovery and phylogenetic reconstruction of Illumina paired-end reads from AIEC isolates were carried out by mapping to the LF82 reference genome (PRJNA33825). Briefly reads were aligned to the LF82 reference with BWA-MEM, variants were called with FreeBayes v1.3.7, SNPs were filtered by base quality (Q ≥ 20) and mapping quality (MAPQ ≥ 30), and variant annotation was performed with SnpEff 5.1^30^. A core SNP alignment spanning the isolates was generated with snippy-core, yielding both a full multiple-sequence alignment and a SNP-only alignment. Putative recombinant regions were removed by applying Gubbins v2.4.1^31^ to the full alignment. A maximum-likelihood phylogeny was inferred from the recombination-filtered core alignment using FastTree2.1.10 under the GTR+Γ substitution model, resulting in the final phylogenetic tree. Tree support values were calculated by FastTree’s internal Shimodaira–Hasegawa test.

A maximum⍰likelihood phylogeny was reconstructed from the recombination⍰filtered core⍰genome alignment with IQ⍰TREE□2.2.6. Such analysis employed the GTR□+□Γ nucleotide⍰substitution model. Node reliability was assessed by performing 1.000 ultrafast bootstrap (UFBoot2) replicates. The resulting tree was rooted on the designated reference strain. Finally, the phylogeny and associated isolate metadata were visualized and explored interactively using Microreact^32^.

### Invasiveness capacity of AIEC bacteria

AIEC strains were analyzed *in vitro* for their abilities to invade intestine-407 epithelial cells (ATCC, CCL-6) by conducting gentamicin protection assays, as previously described^6^. The results were expressed as the number in CFU/mL of invasive bacteria recovered in cells after a 3-hours period of AIEC infection at a MOI of 10 followed by 1 h of gentamicin exposure (100 μg/ml). The assays were performed in triplicates using the non-AIEC *E. coli* strain K-12, as the negative control, and the AIEC reference strain LF82 as the positive control.

### Biofilm formation assay

AIEC strains were cultured overnight in LB medium at 37 °C. Overnight cultures were diluted 1:200 in LB, then the cultures were incubated in flat-bottom 96-well polystyrene plates (Nunc) for 24□h at 37□°C to allow for biofilm formation. At the end of the incubation, growth was measured by reading the OD at 600 nm on a SpectraMax ABS reader. Wells were then washed twice with deionized water to remove residual medium and non-adherent bacteria. Attached biofilms were stained with 0.1% (w/v) crystal violet (Sigma-Aldrich) for 10 min at room temperature, followed by two additional washes with deionized water to eliminate excess dye. Crystal violet-stained biofilms were solubilized in 95% (v/v) ethanol and absorbance was measured at 595□nm using the SpectraMax ABS Reader. OD_595nm_ readings were divided by the optical density of the cultures to obtain normalized biofilm production.

### Motility assay

Bacteria were grown overnight at 37 °C in LB. Cultures were then normalized to an OD of 1 at 600 nm and 1 μl was spotted onto LB agar (0.25 % w/v) and incubated for 7 h. After incubation, plates were imaged using a ChemiDoc XRS+ system (Bio-Rad), and the diameter of the motility zone was measured from the digital images using ImageJ software.

### Quantification of fecal bioactive load of flagellin and lipopolysaccharide

Flagellin and lipopolysaccharide (LPS) were quantified as previously described^33^ using human embryonic kidney (HEK)-Blue-mTLR5 and HEK-Blue-mTLR4 cells, respectively (Invivogen, San Diego, California, USA). Bacteria were grown overnight at 37 °C in LB. Bacteria were subcultured in LB with or without polysorbate80 (1%). The following day, growth was measured by reading the OD at 600 nm on a SpectraMax ABS reader. Samples were then centrifuged at 13.000g for 5 min and the resulting supernatants were serially diluted and applied to mammalian cells. Purified *E. coli* flagellin and LPS (Sigma, St Louis, Missouri, USA) were used for standard curve determination using HEK-Blue-mTLR5 and HEK-Blue-mTLR4 cells, respectively. After 24 hours of stimulation, cell culture supernatants were applied to QUANTI-Blue medium (Invivogen, San Diego, California, USA) and measured alkaline phosphatase activity at 620 nm.

### Prediction of flagellin production based on clone genomes

The association between mutations, virulence traits and flagellin level after P80 exposition was evaluated. Receiver operating characteristic (ROC) curves were calculated (R version 4.1.2, randomForest 4.7-1.1 package, ROCR package) using training data set and validation data set containing randomly affected 70% and 30% of strain, respectively. ROC calculation was repeated 50 times with random sampling of the training and validation data and area under curve (AUC) measurement for each iteration. Mean AUC and standard deviation are presented for each graph.

### Competitive index measurement

For competitive experiments, one strain was selected per condition was selected, and we performed competition *in vitro* and *in vivo* using a 50/50 mix of water-evolved strain and emulsifier-evolved strain. To distinguish each member of these pairs, one member was tagged with a kanamycin-resistance marker carried on a low-copy plasmid. Of note, such tagging was not impacting nor growth nor fitness (**Figure S6A-C**).

#### *in vitro* competitive index measurement

AIEC evolved strains of interest were grown overnight under antibiotic selection (100 μg/ml ampicillin or 100 μg/ml ampicillin + 50μg/ml kanamycin). Bacterial cultures were diluted 50-fold in LB broth, grown overnight at 37□°C without shaking, and both strains were then inoculated in LB with a 1:1 ratio. The obtained inoculum was plated on Drigaslki agar plate with selective antibiotics in order to obtain the input ratio between the two strains. Co-cultures were then sub-cultured twice per day for 3 days, and at each time point, the obtained co-culture was diluted and plated onto Drigalski agar plates with selective antibiotics to determine the CFU per strain, and competitive index was calculated as follow (**Figure 3A**):

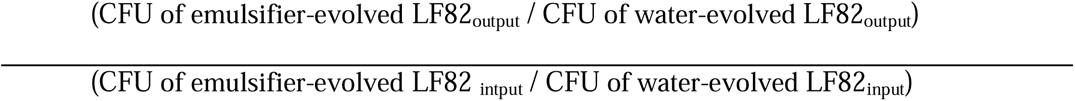

**Figure 3.**
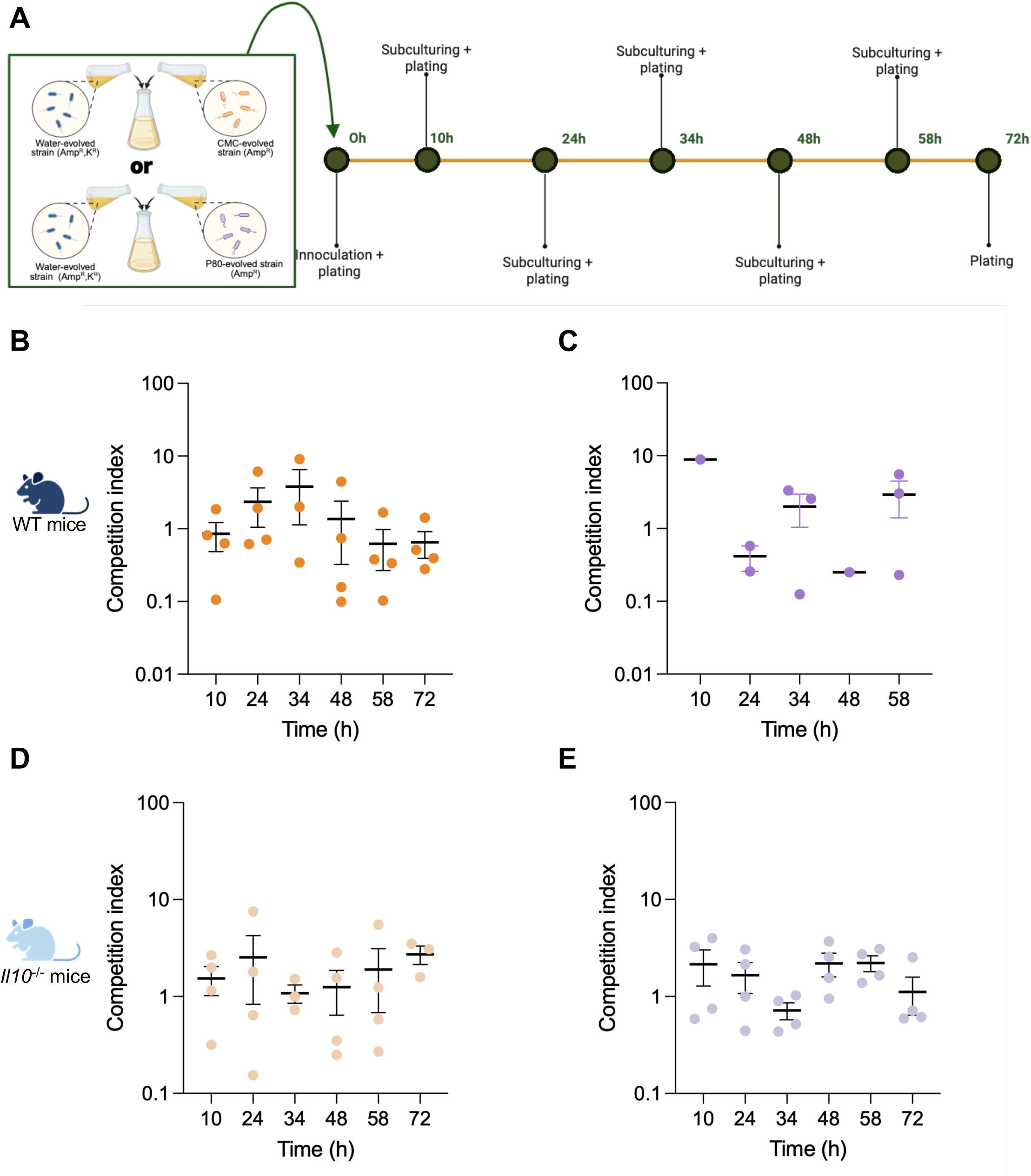
Dietary emulsifiers-driven genomic evolution of AIEC does not alter bacterial *in vitro* growth nor fitness. (**A**) Experimental design of the *in vitro* competition assay. Additive-evolved and water-evolved isolates were mixed 1:1 in LB, passaged twice daily for three days, and plated on selective Drigalski agar at each passage. (**B-C**) CI of WT recovered CMC (**B**) and P80 (**C**) -strains relative to its water-evolved control. (**D-E**) CI of *Il10*^-/-^ recovered CMC (**D**) and P80 (**E**) -evolved strain relative to its water-evolved control. Data are shown as average□±□SEM; individual symbols denote independent replicates. CI: competition index.

#### *in vivo* competitive index measurement

Four weeks old male C57BL/6 mice were purchased from Janvier Laboratories. Animals were housed in a specific pathogen-free barrier facility. Four to 8 mice per group were pre-treated with 25 mg of streptomycin 24 h prior to infection with a 1:1 ratio (5×10^8^ CFU each) of the strains of interest. Briefly, *E*. *coli* strains were grown overnight under selection (100 μg/ml ampicillin or 100 μg/ml ampicillin + 50μg/ml kanamycin). The bacterial cultures were diluted 50-fold in LB broth and grown overnight at 37□°C without shaking. The inoculum was plated on Drigaslki agar plate with selective antibiotics to obtain the input ratio between the two strains. Following infection, the number of bacteria was monitored in fecal output using Drigalski plates with appropriate antibiotic, and the antibiotic resistance profiles of these strains were used to differentiate strains. Six days post-infection, animals were euthanized, and the ileum, colon, caecum, and mucus were homogenized in sterile PBS and then plated on selective media to obtain the colony forming unit (CFU) per strain. The competitive index was calculated as follow (**Figure 4A**):

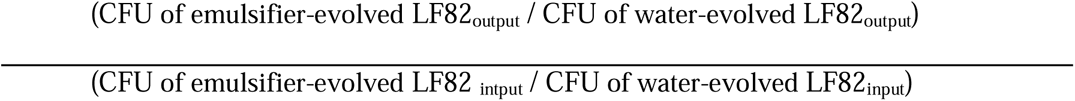

**Figure 4.**
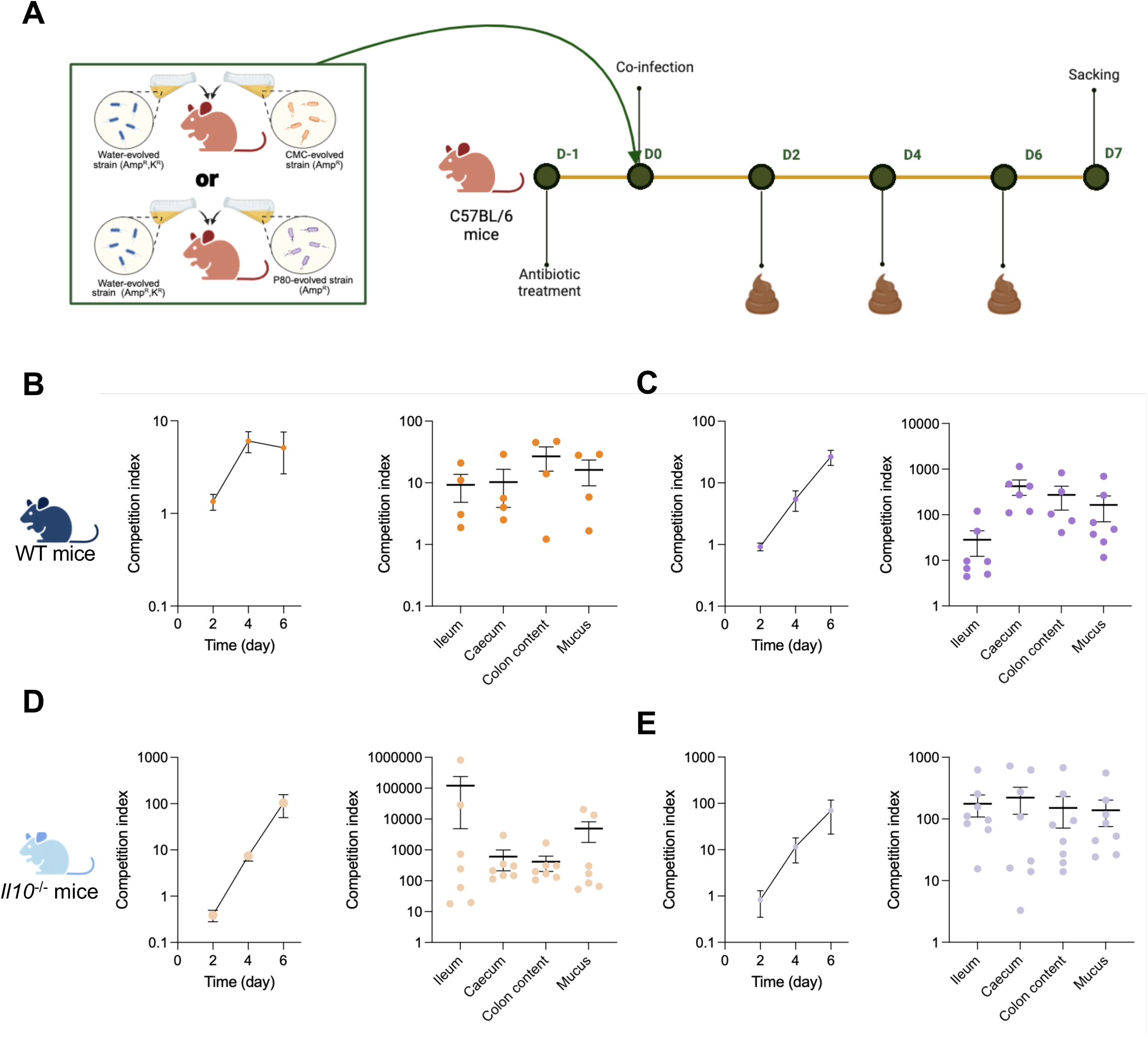
Dietary emulsifiers-driven genomic evolution of AIEC increase their intestinal colonization capacity. (**A**) Overview of the experimental plan used. Streptomycin-pretreated mice received a 1:1 gavage (10^9^ CFU total) of an emulsifier-evolved strain and its water-evolved counterpart. Fecal CFUs were enumerated on days 2, 4 and 6; on day 7, ileum, caecum, colonic contents and mucus were harvested to compute the CI. (**B-C**) CI of CMC- (**B**) and P80- (**C**) evolved strains from WT mice. (**D-E**) CI of CMC- (**D**) and P80- (**E**) evolved strains from *Il10*_*/*_ mice. Data are shown as average□±□SEM; individual symbols represent single animals.

### Statistical analysis

When normality and homoscedasticity postulates were valid, significance was tested using two-way group analysis of variance (ANOVA) with Sidak’s multiple comparisons test (GraphPad Prism software, version 10). When normality of the data was not observed, statistical significance was assessed by the Kruskal Wallis test. Corrected *p*-values are indicated on plots. For phylogenetic and clustering analyses, Hamming distances were calculated. PERMANOVA (adonis2, vegan) and the Association Index, each with 9.999 permutations, were performed to test whether isolates clustered by treatment (water, CMC, P80). To quantify parallel evolution, the number of mutations that were present in at least□2 independent clones (recurrent mutations) were counted for each treatment and compared the proportion of recurrent mutations to the water control by Fisher’s test.

## Results

### Host inflammatory status and dietary emulsifiers independently and synergistically shape AIEC evolution

Diet profoundly influences the behavior and adaptation of intestinal bacteria, including pathobionts such as AIEC that are linked to chronic inflammatory conditions^22,34,35^. To explore the evolutionary impact of chronic dietary emulsifier exposure in the presence or absence of inflammation, we monocolonized wild-type (WT) and *Il10*^-/-^ mice with the AIEC reference strain LF82. Since the route of administration does not qualitatively alter the inflammatory outcome of emulsifier in these models, mice were exposed to CMC, P80, or plain water in drinking water over a three⍰month period (**Fig. 1A**)^36,37^. Four AIEC isolates per mouse were randomly collected and analyzed by whole-genome sequencing.

Such analysis revealed marked differences in the mutational landscape driven by both host genotype and treatment (**Fig. S1, S2A-B, Table 1**). In WT mice, a total of 58 mutated genes were identified: 15 in water-evolved isolates, and 15 and 28 in P80- and CMC-evolved isolates respectively. In *Il10*^-/-^ mice, mutated genes count was substantially higher, with 90 in water-evolved, 20 in CMC-evolved, and 61 in P80-evolved strains. Moreover, emulsifier exposure influenced not only the frequency but also the spectrum of mutations. Both CMC and P80 reduced the occurrence of deletions per strain relative to the water-evolved group in WT strains, whereas CMC increased the proportion of substitutions in WT strains and P80 increased substitution in *Il10*^-/-^ isolates (**Fig. S2C, D**). Remarkably, both emulsifiers significantly reduced the mutation rate per clone while concurrently P80 induced more non-synonymous mutations than synonymous mutation per strain in WT mice (**Fig. S2E, Table 1**). Of note, additional mutations could exist in clones that were not isolated for sequencing. We finally observed that CMC had a similar effect in *Il10*^-/-^ mice (**Fig. S2F**). Analysis of mutations revealed that *Il10*^-/-^ isolates from water- or P80-exposed almost all harbored mutations in the *mutL* gene, which is involved in DNA repair, potentially explaining their higher mutation rate (**Fig. S3**).

**Table 1:**
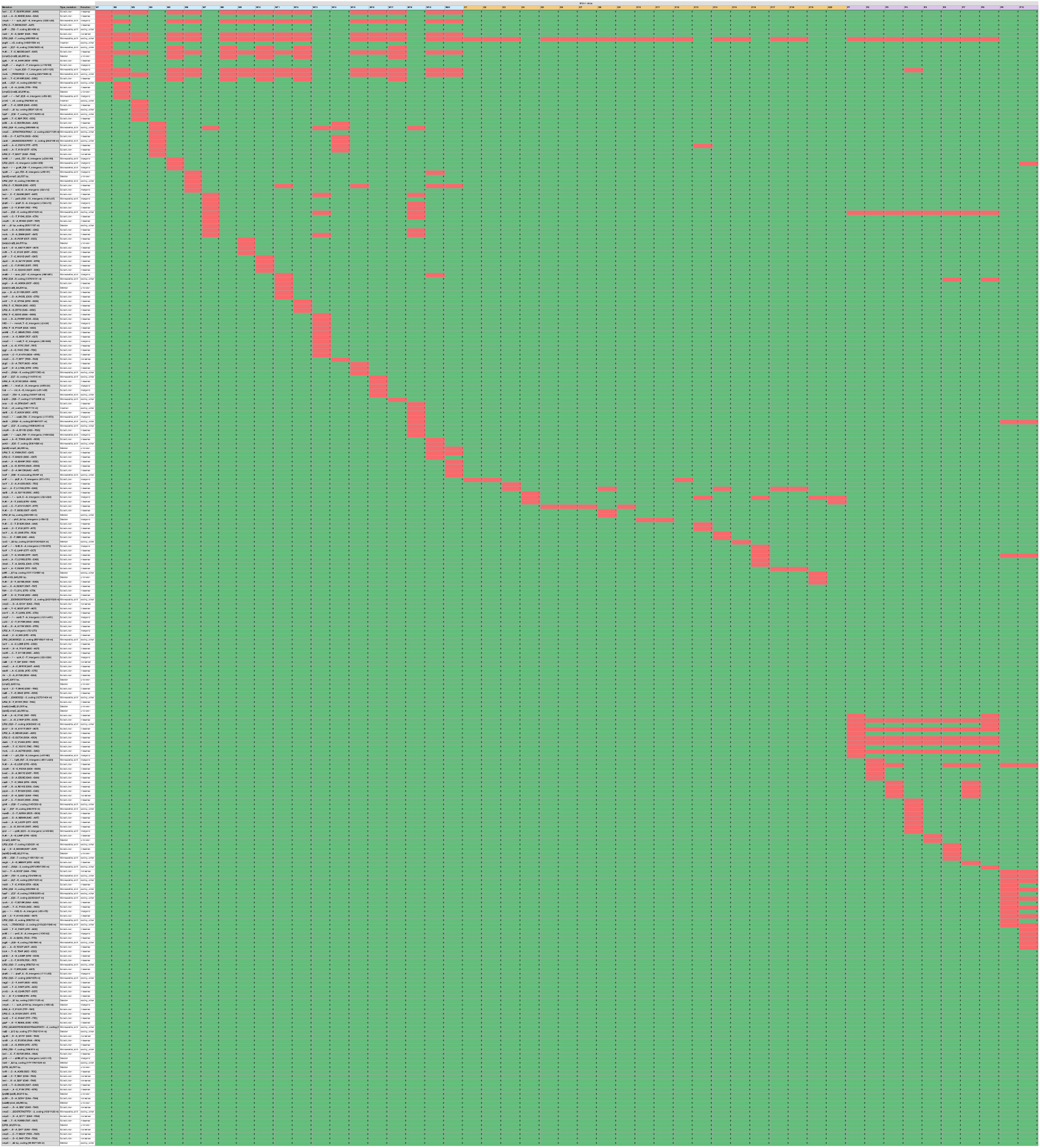
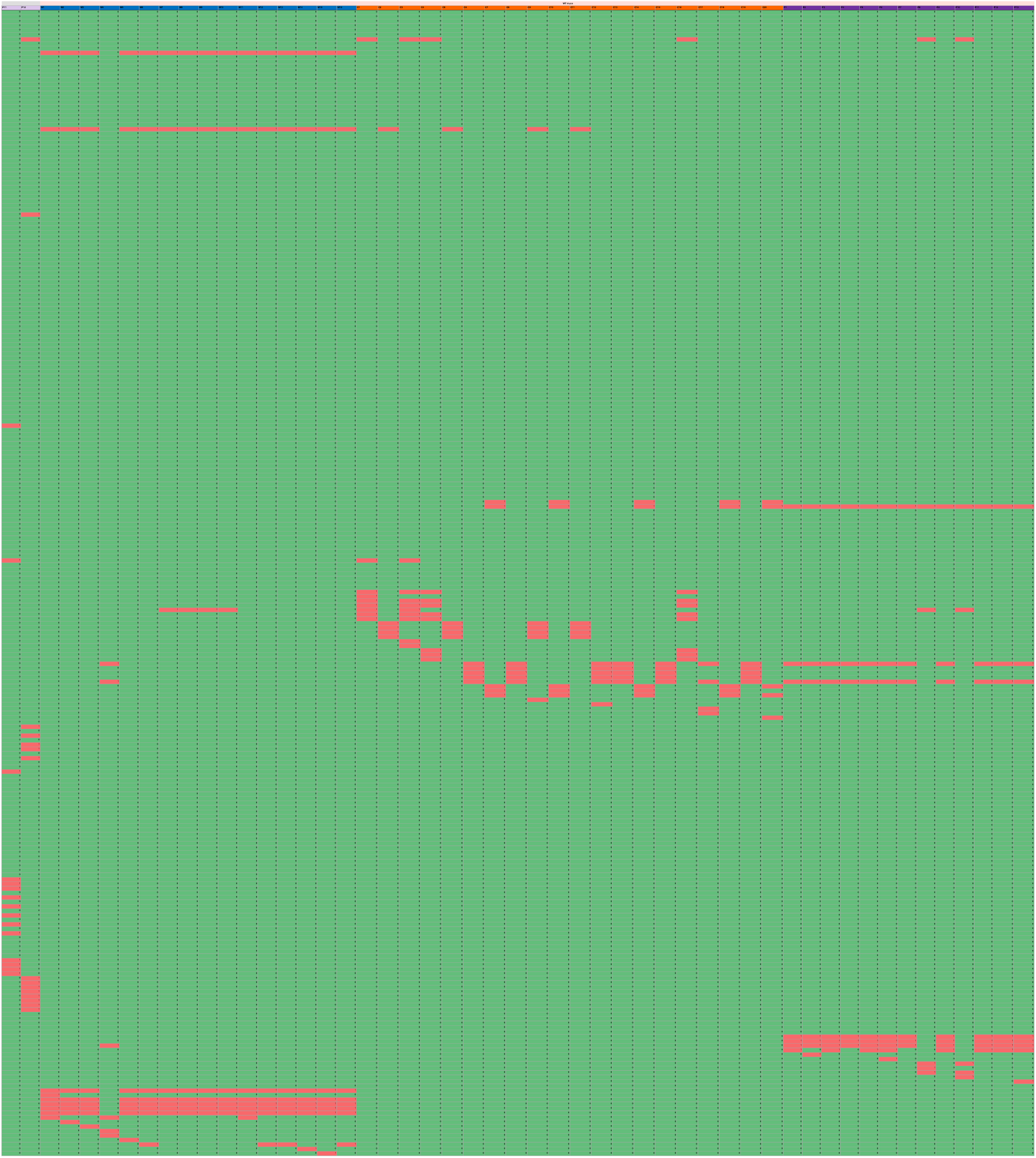
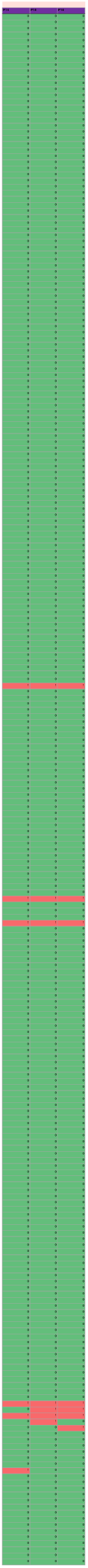
Mutational profiles of evolved strains. The dataset includes the total number of mutations identified in each isolate.

Phylogenetic reconstruction importantly underscored these patterns: isolates segregated into distinct clusters according to treatment in both genotypes (**Fig. 1B, C**). PERMANOVA confirmed a highly significant treatment effect for both genotypes (WT p□=□0.0001; *Il10*⁻/⁻ p□=□0.0001). The Association Index likewise indicated strong non⍰random association between treatment and tree topology (WT AI□=□0.00957, p□=□0.0001; *Il10*⁻*/*⁻ AI□=□0.020, p□=□0.0001). In WT mice, CMC exposure enhances convergent evolution (43□% of all observed mutations are shared by at least two isolates), whereas P80 does not alter the baseline convergence seen in water. Conversely, in *Il10^-/-^*mice, CMC suppresses convergence (only□8□% recurrent mutations) relative to water, while P80 again behaves like the control. These data suggest that treatment strongly shapes both overall genetic distances and the degree of mutational convergence, but the direction of the effect depends on the underlying host genotype. Notably, while PCoA based on mutational profiles of individual isolates from WT mice exposed to water or P80 showed some overlapping points, isolates from WT mice exposed to CMC displayed no evidence of clonal amplification (**Fig. 1D**). In contrast, PCoA of isolates from *Il10*^-/-^ mice revealed distinct positions for most samples, consistent with the absence of clonal amplification within conditions, while highlighting treatment-specific clusters (**Fig. 1E**). Moreover, whereas WT-derived strains converged into tight and condition-specific clades, *Il10*^-/-^-derived isolates displayed greater intra-group diversity and mutational burden (**Fig. S4**). This importantly suggests that underlying inflammation creates a more variable evolutionary landscape for AIEC, while emulsifiers promote more parallel, additive-dependent evolutionary trajectories.

### Emulsifier-driven evolution enhances AIEC pathogenic traits in vitro

Having established emulsifier-driven genomic diversification, we next assessed the functional consequences of accumulated mutations. For this purpose, all recovered isolates were systematically tested for key virulence traits: motility, epithelial invasiveness, biofilm formation, and production of major pro-inflammatory molecules (**Fig. 2**). In WT-derived strains, P80-evolved isolates were notably more invasive in epithelial cell assays while CMC-evolved isolates exhibited significantly increased motility compared to water-evolved controls (**Fig. 2A, B**). Moreover, both emulsifier-evolved groups also produced higher levels of bioactive LPS, a major trigger of host inflammation^38^ (**Fig. 2C**). Conversely, in the *Il10*^-/-^ background, CMC-evolved strains displayed enhanced invasiveness and biofilm formation, but decreased LPS activity relative to water-evolved isolates (**Fig. 2E, G, H**). For P80-evolved strains, no consistent virulence enhancement was observed *in vitro*, suggesting that the strong inflammatory environment overrides emulsifier-induced adaptation in some respects. Overall, except for the selective pressure exerted by dietary emulsifiers, water-evolved strains between both genotypes showed largely unaltered virulence phenotypes, with the only notable exception being a modest decrease in biofilm formation in the *Il10*⁻/⁻ lineage (**Fig. S5**). Such phenotypic stability suggest that shifts observed in CMC- and P80-evolved isolates result specifically from emulsifier-driven selection rather than from genetics background-mediated mutational drift.

Given the recognized role played by flagellin in AIEC virulence, mucus colonization and host pro-inflammatory signaling, we next measured bioactive flagellin with a cell-based TLR5 reporter assay^22,39^. Under basal conditions (i.e. no strain restimulation with emulsifier), P80-evolved isolates from WT mice had reduced flagellin, while CMC showed no effect (**Fig. S6A**). Based on prior *in vitro* findings that P80, unlike CMC, increases flagellin levels, and to model a realistic scenario where evolved pathobionts colonize hosts under continuous emulsifier exposure, we next examined whether emulsifier-evolved strains exhibit differential flagellin responses to P80 compared to water-evolved strain^22^. Upon re-exposure to P80, *Il10*^-/-^-derived CMC and P80 strains generated significantly more bioactive flagellin than their water-evolved counterparts (**Fig. S6D**), suggesting a primed response state likely shaped by the host environment and prior emulsifier exposure. To further understand the genetic determinants of these phenotypes, we next used predictive modeling integrating mutational profiles and observed virulence traits. While no single mutation was sufficient to predict flagellin output, mutation combinations unique to each host background robustly identified high-flagellin evolved strains (**Fig. S6E, G**). In WT isolates, this was driven by mutations in *rfe*, *rssB* and *rpoB*; while in *Il10*^-/-^ isolates, this was driven by mutations in *ybcV*, *rpoC*, and *lacI* (**Fig. S6F, H**). These observations importantly support the idea that emulsifier-driven evolution selects mutational patterns enhancing pathobiont adaptation depending on the intestinal environment.

#### Emulsifier-evolved AIEC outcompete ancestors and harbor in vivo fitness advantage

To test whether these evolved traits confer colonization advantage, we next performed direct competition experiments, both *in vitro* and *in vivo*. For each condition (CMC or P80, WT or *Il10*^-/-^), a representative emulsifier-evolved clone was selected based on several criteria, including intermediate genomic divergence and modest increases in motility, epithelial invasion, and pro-inflammatory molecule secretion (**Fig. S7**). The selected clones were tagged with a kanamycin resistance gene and competed against a representative water-evolved strain (**Fig. S7**). Such kanamycin-resistant emulsifier-evolved strains enabled precise tracking, without any impact of resistance markers on fitness, *in vitro* invasion and colonization ability, as presented **Fig. S8A-D**.

In *in vitro* setting, through repeated cultures of an initial 1:1 inoculum combining a water-evolved clone with an emulsifier-evolved clone, importantly revealed that competitive index stably stayed close to 1 through a 3 days period, representing 6 subculturing of the initial inoculum (**Fig. 3)**. Such observation highlights that, *in vitro*, emulsifier-evolved and water-evolved strains, either in a WT or in an *Il10*^-/-^ genetical background, are not significantly differing in their fitness, growth, nor access to nutriment. However, when performing similar longitudinal analysis of competitive index in a streptomycin-pretreated mouse model (**Fig. 4A**), in all four competition pairs tested, emulsifier-evolved isolates rapidly gained strong intestinal dominance compared to its water-evolved counterpart (**Fig. 4B-E**). This was evidenced by competitive index reaching level between 10 and 1000, meaning that emulsifier-evolved strain dominated by 90 to 99,9% the various intestinal niche evaluated (ileal content, caecum content, colonic content, colonic mucus, as well as longitudinally collected feces, **Fig. 4B-E, Fig. S8E-F**). Importantly, the benefit was even more pronounced in AIEC strains isolated from CMC exposed *Il10*^-/-^ mice, underscoring the potential synergy between host inflammation and dietary emulsifiers in selecting more competitive, adapted AIEC variants.

Overall, our results reveal that broadly used dietary emulsifiers act as powerful evolutionary pressures accelerating the emergence of highly adapted and pathogenic AIEC pathobiont within the gastrointestinal tract. These findings provide a mechanistic basis for the observed increase of AIEC in CD patients and raise wider concerns about the potential role of processed food ingredients in shaping evolution of intestinal pathobionts implicated in chronic disease.

## Discussion

Crohn’s disease (CD) exemplifies the complex interplay between genetics, environment, and the intestinal microbiota in the development of chronic inflammatory disorders. While patients display heterogeneous manifestations, a consistent finding is the high prevalence of adherent-invasive *Escherichia coli* (AIEC) in individuals with CD compared to healthy controls^5,6,40^. Intriguingly, AIEC strains do not belong to a single evolutionary lineage but are scattered across *E. coli* phylogroups, suggesting their independent and recurrent emergence through adaptation to disease-specific selective pressures.

Here, we report that commonly used dietary emulsifiers, carboxymethylcellulose (CMC) and polysorbate 80 (P80), can act as potent evolutionary drivers, accelerating the adaptation of AIEC toward increased pathogenicity, which appears exacerbated in the context of chronic host inflammation. Using mono-colonized murine models, we report that exposure to these food additives, alone or combined with an inflammatory genetical background (*Il10* deficiency), promotes pronounced genomic diversification in AIEC, with distinct mutational signatures depending on both the additive and the host context. Although the evolved strains uniformly displayed a competitive advantage *in vivo*, phenotypes observed *in vitro* differed markedly between WT and *Il10^-/-^* isolated strains. In WT⍰derived isolates, both CMC and P80 exposure were associated with increased LPS level, classically linked to heightened pro⍰inflammatory potential. In contrast, in the *Il10^-/-^* background, these emulsifiers reduced LPS bioactivity while still promoting biofilm formation, especially CMC. Such observation can be explained by the trade⍰off between immune activation and immune evasion that operates under severe inflammation. In a highly inflamed *Il10^-/-^* gut, bacterial strains that dampen LPS signaling may induce reduced detection by the host innate immune system, thereby gaining a survival advantage despite a lower intrinsic pro⍰inflammatory capacity. At the same time, the concomitant increase in biofilm formation can protect the bacteria from antimicrobial peptides and oxidative stress, further supporting persistence. Hence, theses data suggest that the selective pressures imposed by a chronic inflammatory environment shape the direction of emulsifier⍰induced evolution, favoring traits that balance immune activation with immune evasion.

Our findings highlight that while inflammation alone drives AIEC evolution, it is the chronic presence of emulsifiers that directs this diversification towards more pro-inflammatory and colonization-competent phenotypes. Hence, the synergy between modern dietary components and host genetics could underpins the rapid emergence and persistence of AIEC strains in the CD population. This aligns with epidemiological evidence associating increased ultra-processed food consumption with rising rates of IBD^4,41^ and provides a plausible mechanism by which rapid cultural changes in diet may translate into new clinical challenges. Although CMC and P80 have been reported to act directly through the intestinal microbiota, other food additives have been reported *in vitro* to impact the host in a microbiota-independent manner^42,43^. Such host effects could also influence the selection pressures on pathobionts and thereby affect their evolutionary paths.

Mechanistically, we observed that specific combinations of mutations, rather than single genetic changes, underlie the heightened flagellin response and virulence observed in food additive-evolved AIEC. In wild-type hosts, mutations in key stress response and cell surface biosynthesis genes such as *rfe* and *rssB* were observed, while *Il10*^-/-^ backgrounds favored other adaptive trajectories. This context-dependent adaptation underscores the dynamic and tailored nature of pathobiont evolution in response to environmental and inflammatory triggers. Furthermore, several mutations were identified in independent clones, suggesting convergent selection, particularly for clones isolated from WT mice. However, a limitation of our study is that samples were collected after long-term colonization, precluding our ability to evaluate the long-term stability of individual mutations. As such, our single time-point analysis likely overlooks transient mutations present at earlier stages, which could be subsequently lost during adaptation to the host environment and emulsifier exposure. Longitudinal studies of AIEC evolution in human cohorts are ongoing and will address this gap while adding relevance to our findings.

While our mono-association models allowed precise evolutionary tracking, the absence of a complex microbiota is a limitation, as intestinal AIEC adaptation inevitably occurs amidst intense microbial competition and functional redundancy^44^. Furthermore, our isolates were obtained from the gut lumen, whereas AIEC’s pathogenic actions are primarily exerted in the mucus-associated niche, a compartment likely subject to different evolutionary pressures. In addition, this study focused on the genetic and phenotypic adaptation of a single specific AIEC strain. Given prior evidence that *E. coli* accumulates more mutations in inflammatory environments, and that Crohn’s disease–associated strains can outcompete those derived from healthy donors, further studies are warranted to determine whether food emulsifiers similarly shape the evolutionary trajectories of other AIEC lineages^45^.

Nevertheless, these results have broad implications, as they suggest that the modern food environment, characterized by widespread additives use, may be driving not only the selection but also the evolutionary acceleration of pro-inflammatory pathobionts such as AIEC. Given that similar additives and processed food constituents are now global dietary staples, it is plausible that comparable mechanisms are at work in the emergence of other microbiota-derived pathobionts implicated in chronic inflammatory diseases beyond CD. Further studies using emulsifier cocktails at dose mimicking real exposure in Crohn’s-disease patients appear needed to determine the contribution of these additives to AIEC evolution and adaptation.

To conclude, this work establishes that select modern food additives act as evolutionary forces that promote the emergence and adaptation of aggressive intestinal pathobionts. These insights urge a reassessment of dietary recommendations for at-risk individuals and highlight the need for further studies on how contemporary diets are shaping the evolution of the human gut microbiota. Understanding these dynamics will be crucial to develop strategies to prevent and manage chronic inflammatory diseases in the modern age.

## Acknowledgment

The authors thank the Genom’IC platforms (INSERM U1016, Paris, France) for their help in library sequencing and Nadia Andrea Andreani for her help and guidance with the phylogenetic analysis. We dedicate this article to Nico Barnich who was a mentor, a colleague and a friend to many of the authors.

## Author contributions

B.C: designed the study; B.C and H.R performed the study; C.C, L.M, J.D contributed to experiments; E.B, O.T, O.E, A.B : contributed to analysis and interpretation of data for the work; E.V provided samples; B.C and H.R.: drafted the manuscript. All authors critically reviewed and approved the final manuscript.

## Data availability

The data underlying this article are available in the article and in its online supplementary material.

## Abbreviations

AIEC: Adherent Invasive *Escherichia coli*
OD: Optical Density
CD: Crohn’s Disease
CMC: Carboxymethylcellulose
LPS: Lipopolysaccharide
P80: Polysorbate 80
WT: Wild-type

## Supplementary figure legends

**Figure S1.**
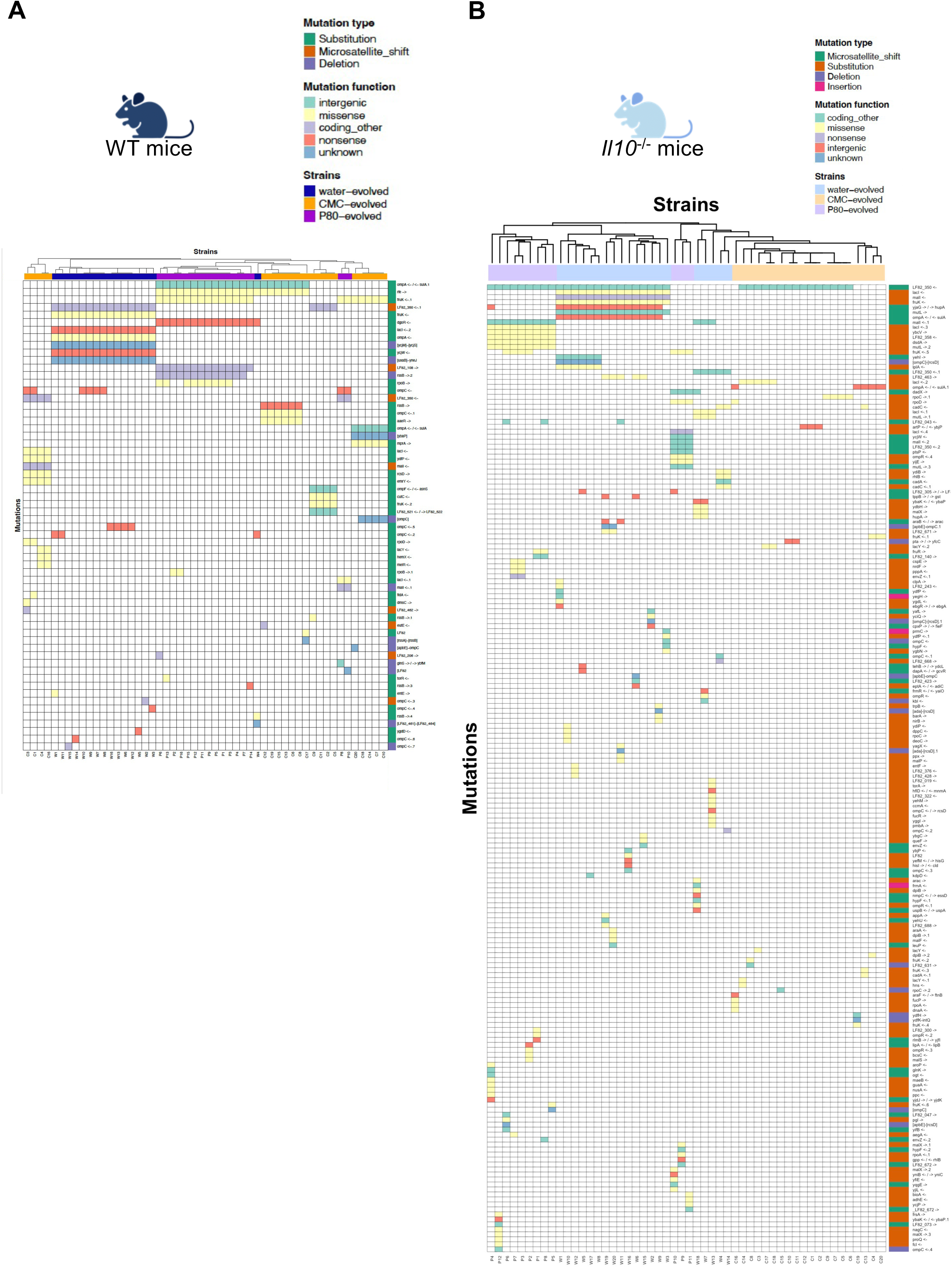
Mutational profile on the evolved strains. Dataset contains total mutated genes for each isolate. Heatmaps of mutated genes detected in WT (**A**) and *Il10*^-/-^ (**B**) -derived strains. Rows correspond to mutated genes and their genomic orientation and columns to individual isolate. Mutation type and function were annotated. Isolates were hierarchically clustered using Bray-Curtis dissimilarity.

**Figure S2.**
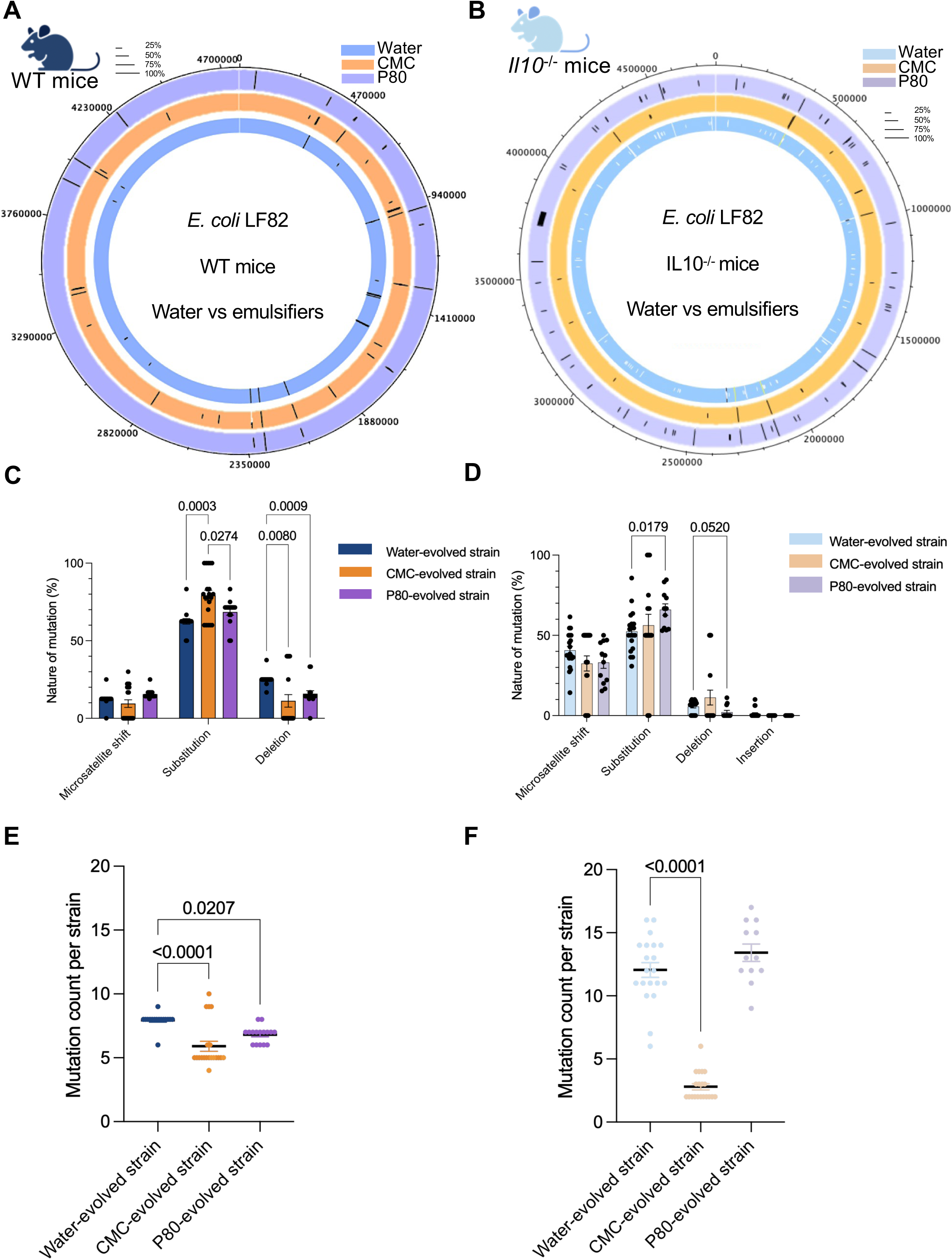
Genotypic characterization of dietary emulsifier-driven AIEC evolution. (**A-B**) Genomic location of mutations identified in WT (**A**) and *Il10*^-/-^ (**B**) -derived strains. Bar length denotes the percentage of mice in which at least one clone carried the indicated mutation. (**C-D**) Proportions of mutation types detected in WT (**C**) and *Il10*^-/-^ (**D**) - derived strains. Data represent mean□±□SEM. ∗p < 0.05 determined by two-way ANOVA test. (**E-F**) Total number of mutations per strain isolated from (**E**) WT mice and (**F**) *Il10*^-/-^mice. Data show median ±□interquartile range. ∗p < 0.05 determined by Kruskall-Wallis test.

**Figure S3.**
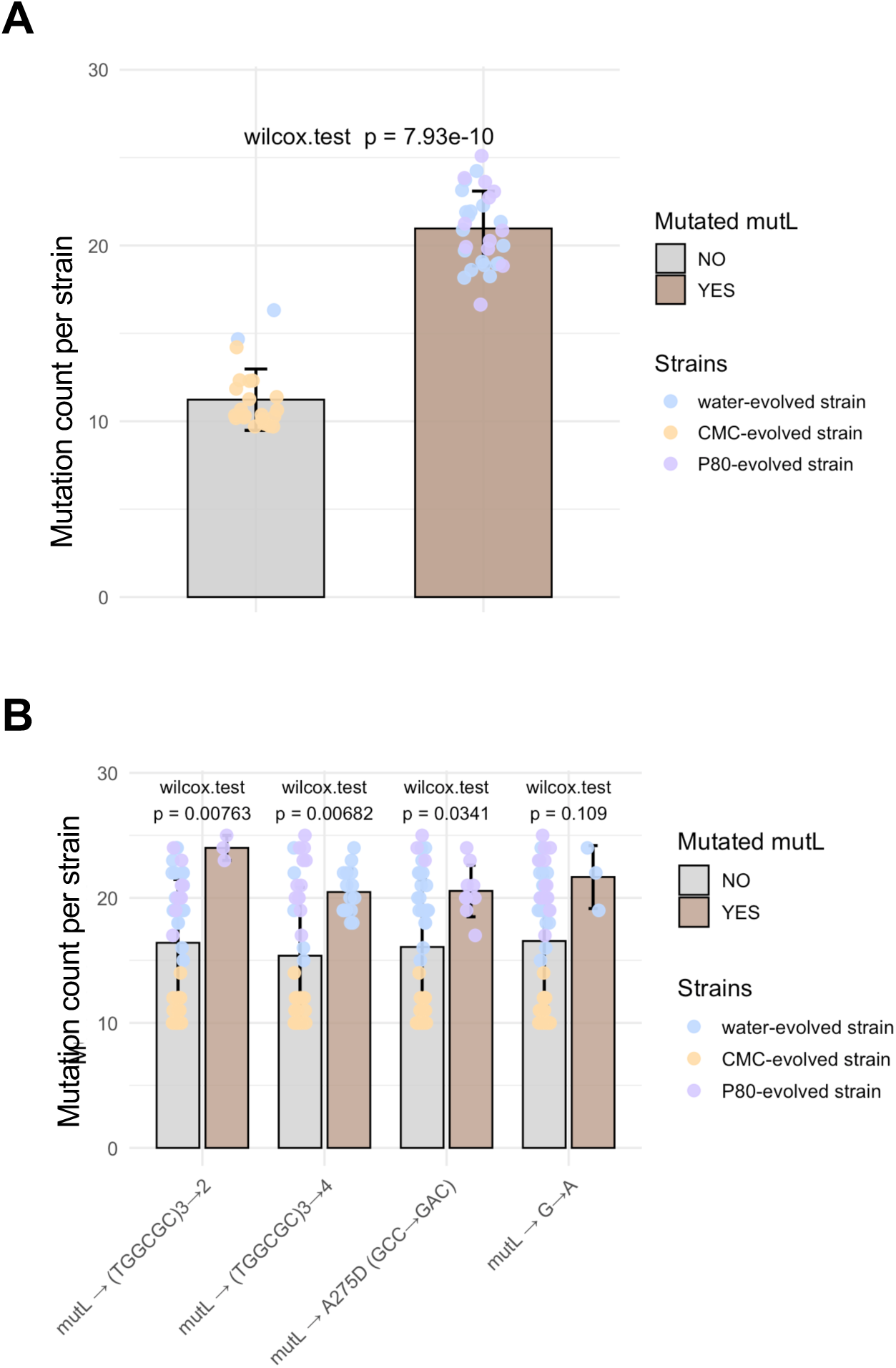
*mutL* mutation influences overall mutation rate. **(A)** Total number of mutations detected in *Il10*^-/-^-derived strains as a function of *mutL* mutation presence. **(B)** Total number of mutations detected in *Il10*^-/-^-derived strains according to *mutL* mutation presence and type. Data represent mean□±□SD. ∗p < 0.05 determined by Wilcoxon test.

**Figure S4.**
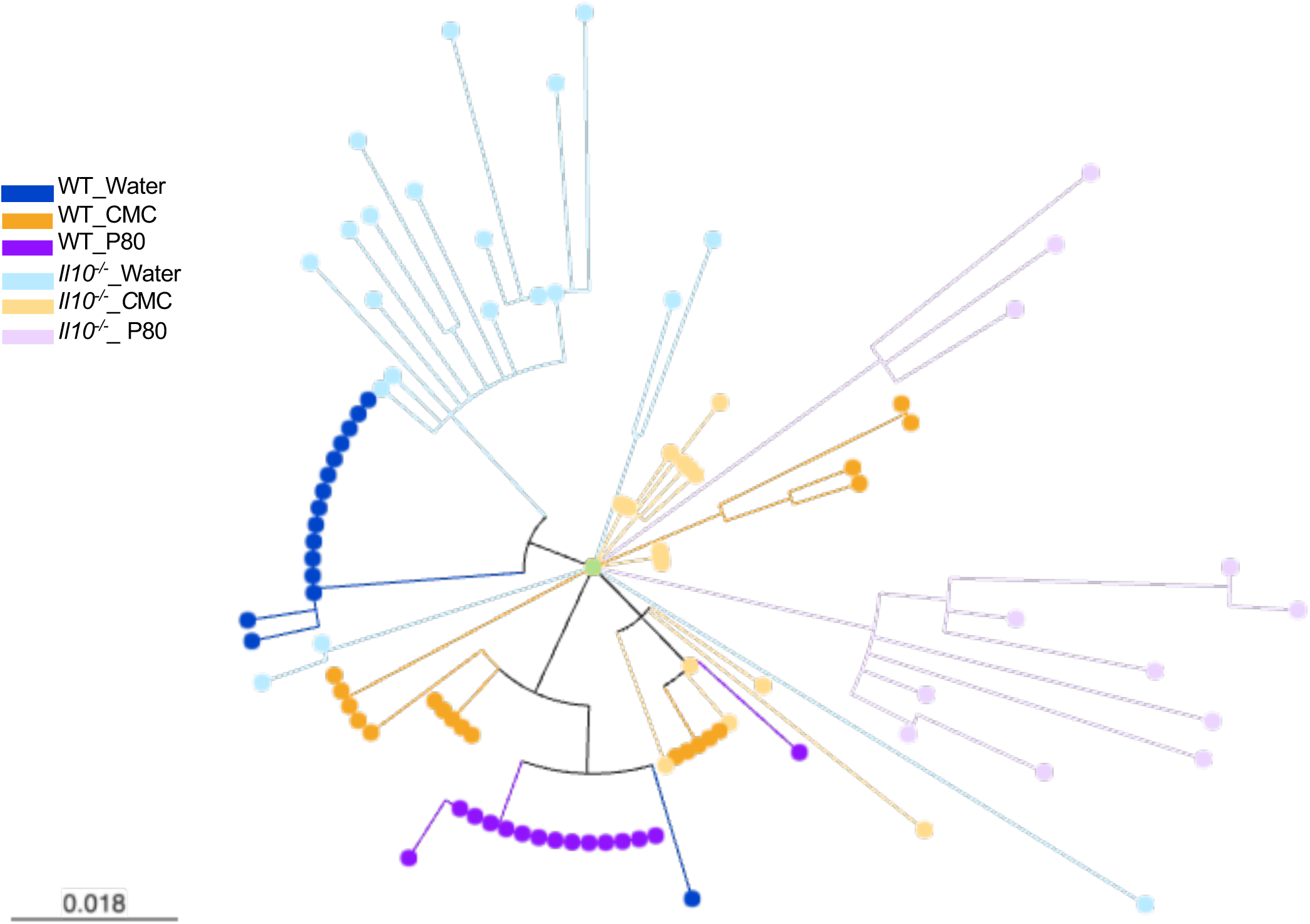
Inflammation and dietary emulsifiers shape AIEC genomic evolution in specific ways. Maximum-likelihood phylogenies of evolved isolates from both WT and *Il10*^-/-^ mice in comparison to the LF82 reference. Branch colors indicate treatment group.

**Figure S5.**
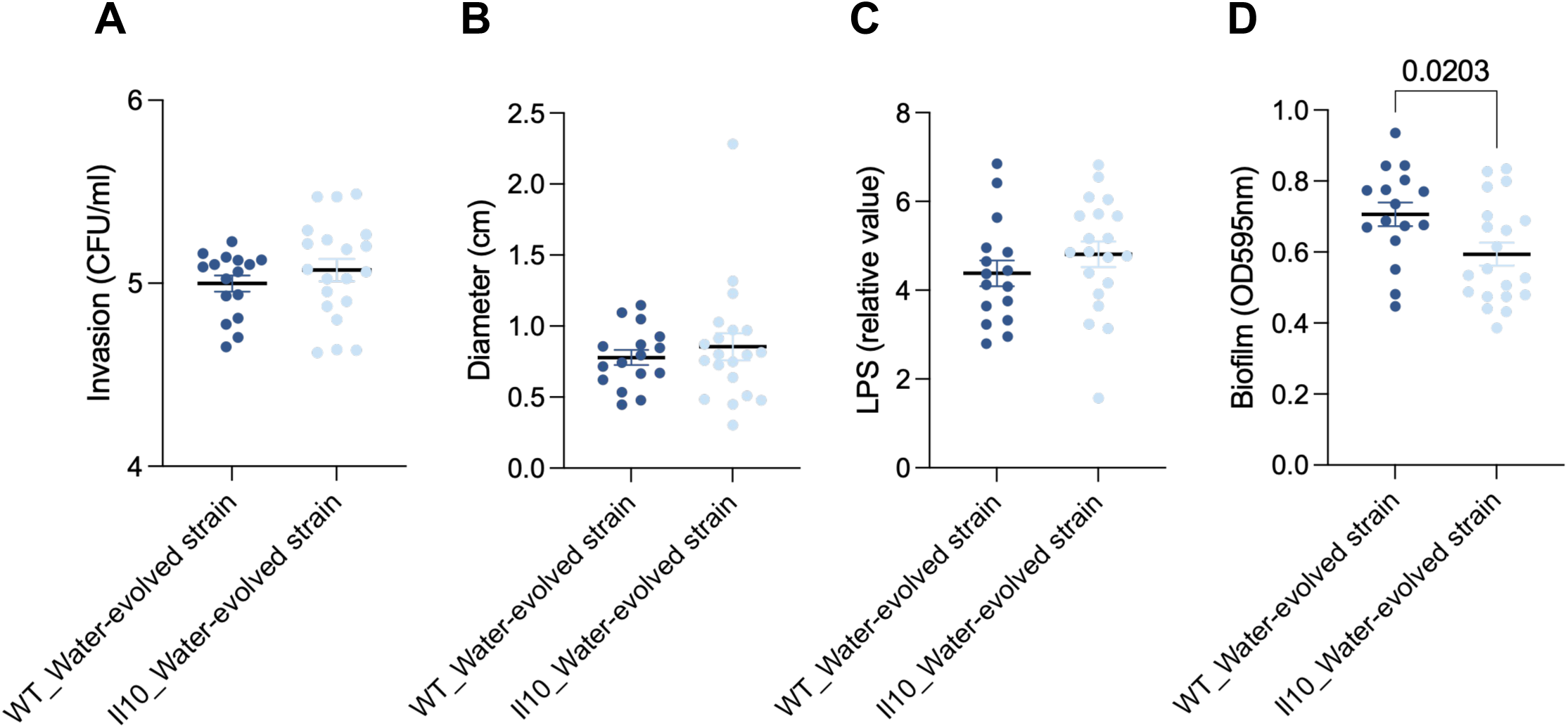
Inflammation-driven genomic evolution of AIEC impacts modestly their pathogenic potential *in vitro*. *in vitro* pathogenic traits of *Il10^-/-^* isolates were assessed by measuring invasion of IECS (**A**) swimming motility (**B**) bioactive LPS level normalized by OD600nm (**C**) and biofilm formation (**D**). Data are shown as the average□±□SEM. ∗p < 0.05 compared to water-isolated strains, determined by t-test.

**Figure S6.**
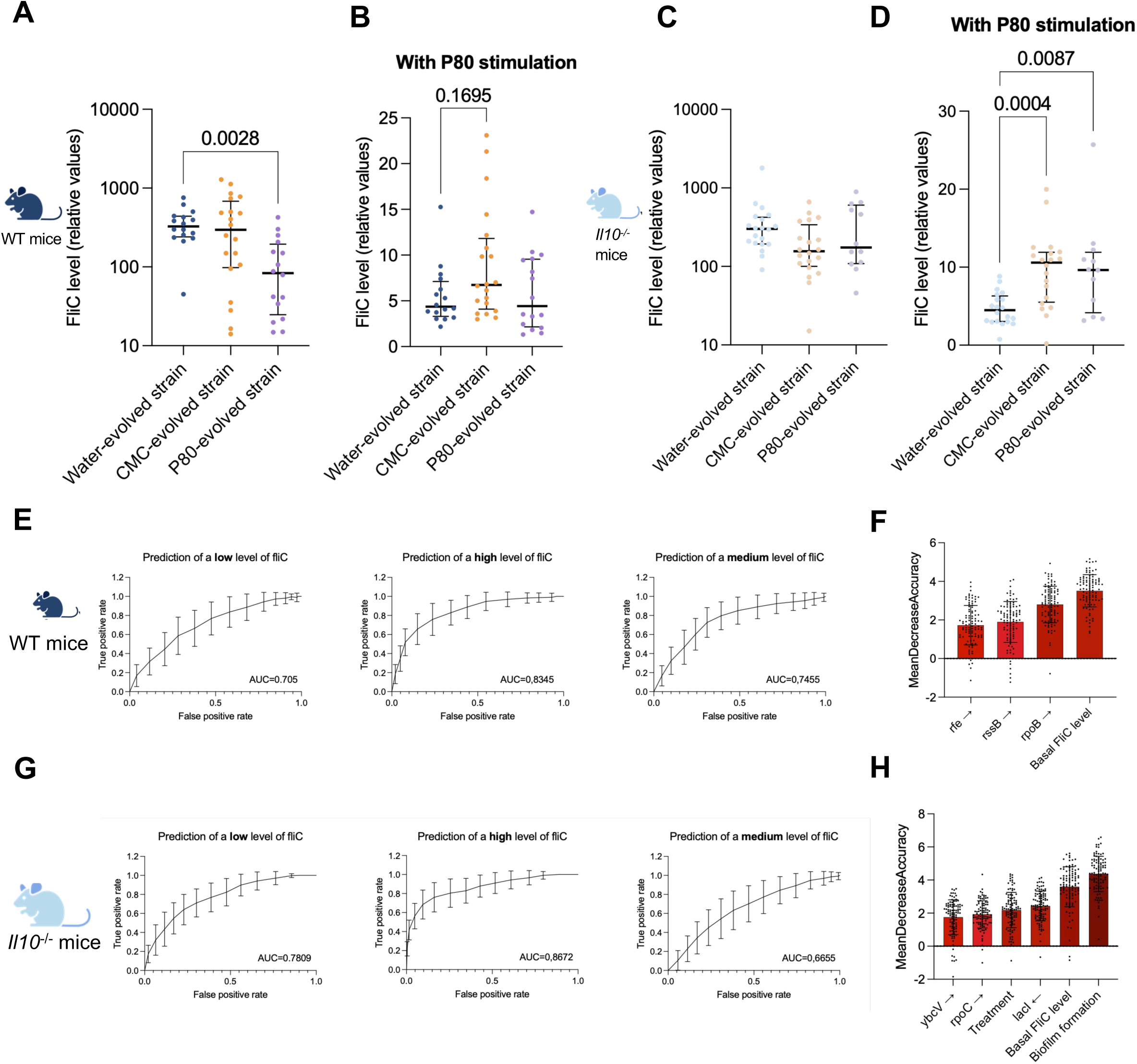
Dietary emulsifier-driven genomic evolution increases AIEC responsiveness to P80, which is predictable from the mutated genes. (**A-D**) Bioactive flagellin levels measured by a HEK-Blue mTLR5 reporter assay in evolved isolates from WT (**A, B**) and Il10⁻/⁻ (**C, D**) mice: (**A, C**) basal secretion, (**B, D**) secretion after stimulation with 1% P80. Data are median□±□interquartile range; *p <□0.05 versus water-evolved strains calculated via Kruskal–Wallis test. (**E, G**) ROC curves (AUC indicated) for random-forest models classifying isolates as high, medium or low flagellin producers post-P80 in WT (**E**) and *Il10*⁻*/*⁻ (**G**) strains (**F, H**) Top driver mutated genes and their orientation ranked by feature-importance in the prediction models for WT (**F**) and *Il10*⁻*/*⁻ (**H**) isolates

**Figure S7.**
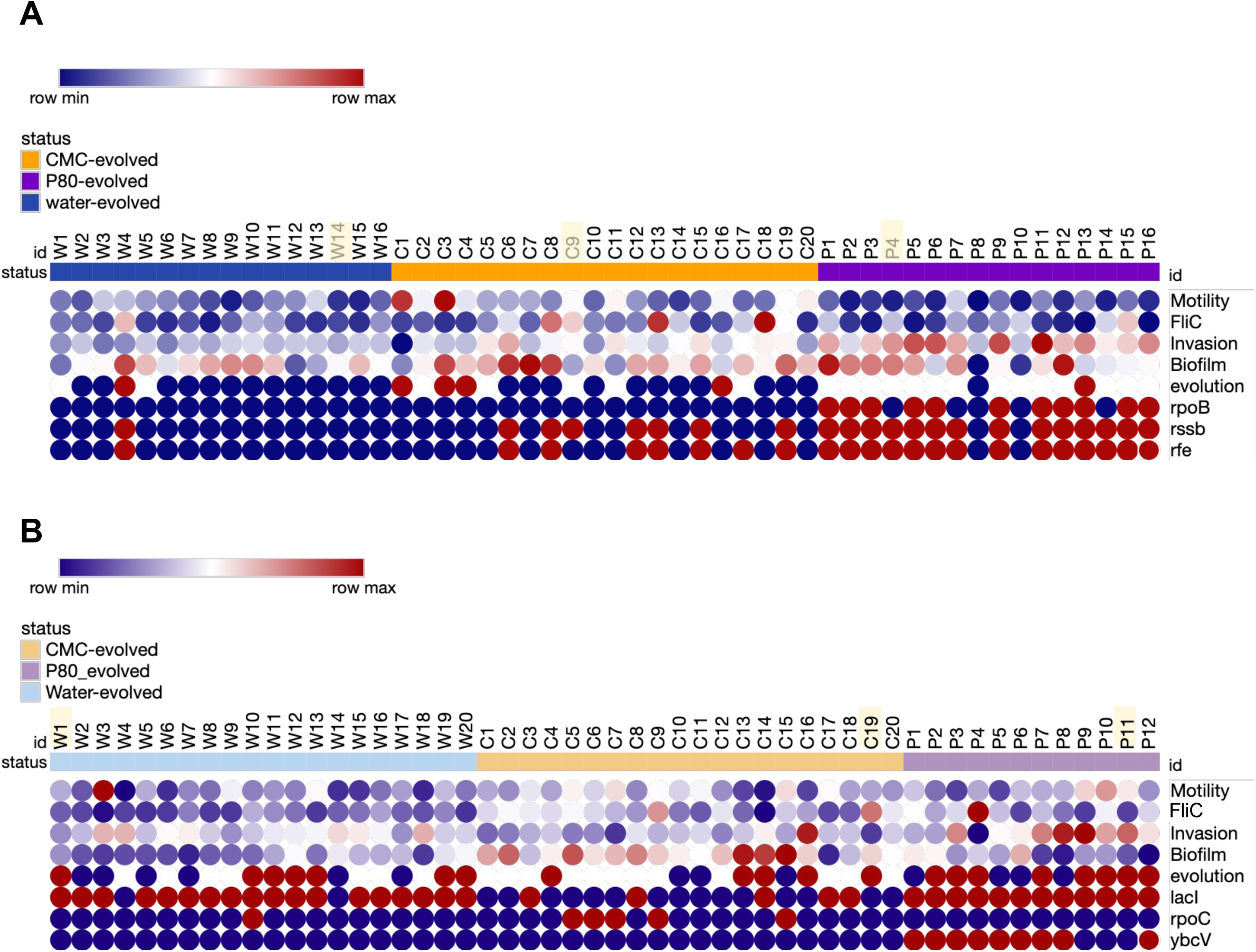
Key characteristics of all isolates. Heatmaps displaying key phenotypic and genotypic features identified in WT (**A**) and *Il10*⁻*/*⁻ (**B**) -derived strains. Rows represent major traits, while columns correspond to individual isolates. Representative strains for each condition are highlighted in yellow.

**Figure S8.**
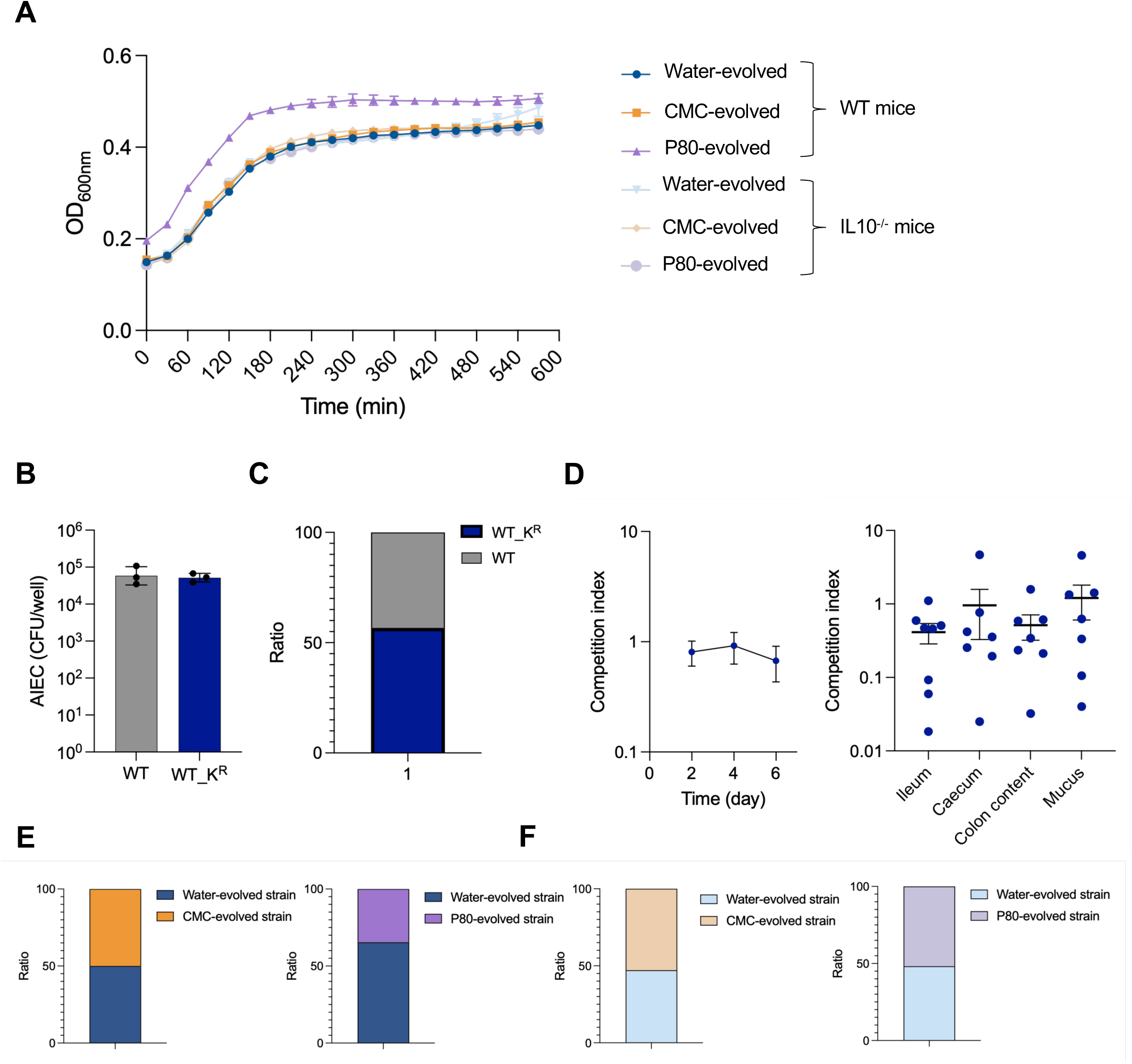
Growth curves and inoculum ratio of bacterial strains used for *in vivo* competition experiments. (**A**) Growth curves of the six representative evolved isolates selected for competition. (**B)** Invasion of intestinal epithelial cells (I407) by parental LF82 and its isogenic kanamycin⍰resistant derivative. Data represent the mean of three independent experiments, each performed in technical triplicate. **(C-D**) Validation of the competition model using parental LF82 and its isogenic kanamycin-resistant derivative. (**C**) Inoculum ratios at gavage (**D**) CI of the kanamycin-resistant strain relative to the control in fecal samples (**left**) and in ileum, caecum, colon content and mucus (**right**) on day□7 post-infection. Data are mean□±□SEM; individual symbols denote single animals. (**E-F**) Gavage inoculum ratios for evolved vs. parental competition pairs recovered from WT (**E**) and *Il10*^-/-^ (**F**) mice

## References

1. Bragazzi, N. L., Del Rio, D., Mayer, E. A. & Mena, P. We Are What, When, And How We Eat: The Evolutionary Impact of Dietary Shifts on Physical and Cognitive Development, Health, and Disease. Advances in Nutrition 15, 100280 (2024).

2. Alt, K. W., Al-Ahmad, A. & Woelber, J. P. Nutrition and Health in Human Evolution–Past to Present. Nutrients 14, 3594 (2022).

3. Cordain, L. et al. Origins and evolution of the Western diet: health implications for the 21st century. Am J Clin Nutr 81, 341–354 (2005).

4. Chen, J. et al. Intake of Ultra-processed Foods Is Associated with an Increased Risk of Crohn’s Disease: A Cross-sectional and Prospective Analysis of 187 154 Participants in the UK Biobank. J Crohns Colitis 17, 535–552 (2023).

5. Darfeuille-Michaud, A. Adherent-invasive Escherichia coli: a putative new E. coli pathotype associated with Crohn’s disease. Int J Med Microbiol 292, 185–193 (2002).

6. Darfeuille-Michaud, A. et al. High prevalence of adherent-invasive Escherichia coli associated with ileal mucosa in Crohn’s disease. Gastroenterology 127, 412–421 (2004).

7. Kaplan, G. G. & Ng, S. C. Understanding and Preventing the Global Increase of Inflammatory Bowel Disease. Gastroenterology 152, 313–321.e2 (2017).

8. Chassaing, B. & Darfeuille-Michaud, A. The commensal microbiota and enteropathogens in the pathogenesis of inflammatory bowel diseases. Gastroenterology 140, 1720–1728 (2011).

9. Kamali Dolatabadi, R., Feizi, A., Halaji, M., Fazeli, H. & Adibi, P. The Prevalence of Adherent-Invasive Escherichia coli and Its Association With Inflammatory Bowel Diseases: A Systematic Review and Meta-Analysis. Front Med (Lausanne) 8, 730243 (2021).

10. Chassaing, B. et al. Crohn disease–associated adherent-invasive E. coli bacteria target mouse and human Peyer’s patches via long polar fimbriae. J Clin Invest 121, 966–975 (2011).

11. Glasser, A. L. et al. Adherent invasive Escherichia coli strains from patients with Crohn’s disease survive and replicate within macrophages without inducing host cell death. Infect Immun 69, 5529–5537 (2001).

12. Rolhion, N., Carvalho, F. A. & Darfeuille-Michaud, A. OmpC and the sigma(E) regulatory pathway are involved in adhesion and invasion of the Crohn’s disease-associated Escherichia coli strain LF82. Mol Microbiol 63, 1684–1700 (2007).

13. Elhenawy, W., Tsai, C. N. & Coombes, B. K. Host-Specific Adaptive Diversification of Crohn’s Disease-Associated Adherent-Invasive Escherichia coli. Cell Host Microbe 25, 301–312.e5 (2019).

14. Unni, R. et al. Evolution of E. coli in a mouse model of inflammatory bowel disease leads to a disease-specific bacterial genotype and trade-offs with clinical relevance. Gut Microbes 15, 2286675 (2023).

15. Adolph, T. E. & Tilg, H. Western diets and chronic diseases. Nat Med 30, 2133–2147 (2024).

16. Malesza, I. J. et al. High-Fat, Western-Style Diet, Systemic Inflammation, and Gut Microbiota: A Narrative Review. Cells 10, 3164 (2021).

17. Roberts, C. L., Rushworth, S. L., Richman, E. & Rhodes, J. M. Hypothesis: Increased consumption of emulsifiers as an explanation for the rising incidence of Crohn’s disease. J Crohns Colitis 7, 338–341 (2013).

18. Chassaing, B. et al. Dietary emulsifiers impact the mouse gut microbiota promoting colitis and metabolic syndrome. Nature 519, 92–96 (2015).

19. Chassaing, B., De Bodt, J., Marzorati, M., Van de Wiele, T. & Gewirtz, A. T. Dietary emulsifiers directly alter human microbiota composition and gene expression ex vivo potentiating intestinal inflammation. Gut 66, 1414–1427 (2017).

20. Chassaing, B. et al. Randomized Controlled-Feeding Study of Dietary Emulsifier Carboxymethylcellulose Reveals Detrimental Impacts on the Gut Microbiota and Metabolome. Gastroenterology 162, 743–756 (2022).

21. Rytter, H. et al. Invitro microbiota model recapitulates and predicts individualised sensitivity to dietary emulsifier. Gut 74, e333925 (2025).

22. Viennois, E. et al. Dietary Emulsifiers Directly Impact Adherent-Invasive E. coli Gene Expression to Drive Chronic Intestinal Inflammation. Cell Rep 33, 108229 (2020).

23. Hecht, G. et al. A simple cage-autonomous method for the maintenance of the barrier status of germ-free mice during experimentation. Lab Anim 48, 292–297 (2014).

24. Choi, K.-H. & Schweizer, H. P. mini-Tn7 insertion in bacteria with single attTn7 sites: example Pseudomonas aeruginosa. Nat Protoc 1, 153–161 (2006).

25. Avilan, L. Assembling Multiple Fragments: The Gibson Assembly. Methods Mol Biol 2633, 45–53 (2023).

26. Ferrières, L. et al. Silent mischief: bacteriophage Mu insertions contaminate products of Escherichia coli random mutagenesis performed using suicidal transposon delivery plasmids mobilized by broad-host-range RP4 conjugative machinery. J Bacteriol 192, 6418–6427 (2010).

27. Martin, M. Cutadapt removes adapter sequences from high-throughput sequencing reads. EMBnet.journal 17, 10–12 (2011).

28. Bolger, A. M., Lohse, M. & Usadel, B. Trimmomatic: a flexible trimmer for Illumina sequence data. Bioinformatics 30, 2114–2120 (2014).

29. Deatherage, D. E. & Barrick, J. E. Identification of Mutations in Laboratory-Evolved Microbes from Next-Generation Sequencing Data Using breseq. in Engineering and Analyzing Multicellular Systems: Methods and Protocols (eds Sun, L. & Shou, W.) 165–188 (Springer, New York, NY, 2014). doi:10.1007/978-1-4939-0554-6_12.

30. Cingolani, P. et al. A program for annotating and predicting the effects of single nucleotide polymorphisms, SnpEff. Fly (Austin) 6, 80–92 (2012).

31. Croucher, N. J. et al. Rapid phylogenetic analysis of large samples of recombinant bacterial whole genome sequences using Gubbins. Nucleic Acids Res 43, e15 (2015).

32. Argimón, S. et al. Microreact: visualizing and sharing data for genomic epidemiology and phylogeography. Microb Genom 2, e000093 (2016).

33. Chassaing, B., Koren, O., Carvalho, F. A., Ley, R. E. & Gewirtz, A. T. AIEC Pathobiont Instigates Chronic colitis in Susceptible Hosts by Altering Microbiota composition. Gut 63, 1069–1080 (2014).

34. Kuffa, P. et al. Fiber-deficient diet inhibits colitis through the regulation of the niche and metabolism of a gut pathobiont. Cell Host Microbe 31, 2007–2022.e12 (2023).

35. Rytter, H., Sturgeon, H. & Chassaing, B. Diet-pathobiont interplay in health and inflammatory bowel disease. Trends Microbiol S0966–842X(25)00112-X (2025) doi:10.1016/j.tim.2025.04.003.

36. Miquel, S. et al. Complete Genome Sequence of Crohn’s Disease-Associated Adherent-Invasive E. coli Strain LF82. PLOS ONE 5, e12714 (2010).

37. Chassaing, B. et al. Dietary emulsifiers impact the mouse gut microbiota promoting colitis and metabolic syndrome. Nature 519, 92–96 (2015).

38. Vijay-Kumar, M. et al. Bacterial flagellin is a dominant, stable innate immune activator in the gastrointestinal contents of mice and rats. Gut Microbes 15, 2185031 (2023).

39. Sevrin, G. et al. Adaptation of adherent-invasive E. coli to gut environment: Impact on flagellum expression and bacterial colonization ability. Gut Microbes 11, 364–380 (2020).

40. Kamali Dolatabadi, R., Feizi, A., Halaji, M., Fazeli, H. & Adibi, P. The Prevalence of Adherent-Invasive Escherichia coli and Its Association With Inflammatory Bowel Diseases: A Systematic Review and Meta-Analysis. Front Med (Lausanne) 8, 730243 (2021).

41. Narula, N. et al. Association of ultra-processed food intake with risk of inflammatory bowel disease: prospective cohort study. BMJ 374, n1554 (2021).

42. Laudisi, F. et al. The Food Additive Maltodextrin Promotes Endoplasmic Reticulum Stress-Driven Mucus Depletion and Exacerbates Intestinal Inflammation. Cell Mol Gastroenterol Hepatol 7, 457–473 (2019).

43. Jiang, H.-Y., Wang, F., Chen, H.-M. & Yan, X.-J. κ-carrageenan induces the disruption of intestinal epithelial Caco-2 monolayers by promoting the interaction between intestinal epithelial cells and immune cells. Molecular Medicine Reports 8, 1635–1642 (2013).

44. Van den Abbeele, P., et al. Arabinoxylans, inulin and Lactobacillus reuteri 1063 repress the adherent-invasive Escherichia coli from mucus in a mucosa-comprising gut model. npj Biofilms Microbiomes 2, 16016 (2016).

45. Ameline, C. et al. Evolution of Escherichia coli strains under competent or compromised adaptive immunity. PLoS Pathog 21, e1012442 (2025).

